# Decreasing ganglioside synthesis delays motor and cognitive symptom onset in *Spg11* knockout mice

**DOI:** 10.1101/2024.01.29.577736

**Authors:** Manon Fortier, Margaux Cauhapé, Suzie Buono, Julien Becker, Alexia Menuet, Julien Branchu, Ivana Ricca, Serena Mero, Karim Dorgham, Khalid-Hamid El Hachimi, Kostantin Dobrenis, Benoit Colsch, Dominic Samaroo, Morgan Devaux, Alexandra Durr, Giovanni Stevanin, Filippo M. Santorelli, Sophie Colombo, Belinda Cowling, Frédéric Darios

**Affiliations:** Sorbonne Université, Paris Brain Institute (ICM Institut du Cerveau), INSERM U1127, CNRS UMR 7225, Assistance Publique-Hôpitaux de Paris (AP-HP), Paris, France; Dynacure SA (now Flamingo Therapeutics NV), Illkirch, France; Molecular Medicine, IRCCS Fondazione Stella Maris, 56128 Pisa, Italy; Sorbonne Université, INSERM, Centre d’Immunologie et des Maladies Infectieuses-Paris (CIMI-Paris), F-75013 Paris, France; EPHE, PSL Research University, Paris, France; Dominick P. Purpura Department of Neuroscience, Albert Einstein College of Medicine, Bronx, NY 10461, USA; Université Paris-Saclay, CEA, INRAE, Département Médicaments et Technologies pour la Santé, MetaboHUB, Gif sur Yvette, France; University of Bordeaux, CNRS, INCIA, UMR 5287, NRGen Team, Bordeaux, France

**Keywords:** Lysosome, ganglioside, neurodegeneration, therapy, genetic disease

## Abstract

Biallelic variants in the *SPG11* gene account for the most common form of autosomal recessive hereditary spastic paraplegia characterized by motor and cognitive impairment, with currently no therapeutic option. We previously observed in a *Spg11* knockout mouse that neurodegeneration is associated with accumulation of gangliosides in lysosomes. To test whether a substrate reduction therapy could be a therapeutic option, we downregulated the key enzyme involved in ganglioside biosynthesis using an AAV-PHP.eB viral vector expressing a miRNA targeting *St3gal5*. Downregulation of *St3gal5* in *Spg11* knockout mice prevented the accumulation of gangliosides, delayed the onset of motor and cognitive symptoms, and prevented the upregulation of serum levels of neurofilament light chain, a biomarker widely used in neurodegenerative diseases. Importantly, similar results were observed upon treatment of *Spg11* knockout mice with venglustat, a pharmacological inhibitor of glucosylceramide synthase expected to decrease ganglioside synthesis. Downregulation of *St3gal5* or venglustat treatment of *Spg11* knockout mice strongly decreased the formation of axonal spheroids, previously associated with impaired trafficking. Venglustat had similar effect on cultured human SPG11 neurons. In conclusion, this work identifies the first disease-modifying therapeutic strategy in SPG11, and provides data supporting its relevance for therapeutic testing in SPG11 patients.

## Introduction

Hereditary spastic paraplegias (HSP) are a group of neurodegenerative diseases characterized by leg spasticity due to progressive degeneration of the corticospinal tract. This group of diseases is genetically highly heterogeneous, with more than 60 genes and 80 loci associated with HSP so far (Darios et al., 2022). Mutations in the *SPG11* gene are responsible for a severe form of autosomal recessive HSP, with patients often presenting upper motor neuron involvement with spasticity, as well as lower motor neuron changes with peripheral neuropathy and other clinical signs such as cerebellar ataxia and cognitive impairment (Hehr et al., 2007; Stevanin et al., 2008). This disease is due to loss of function mutations in the *SPG11* gene, leading to the absence of *SPG11* gene product, spatacsin (Stevanin et al., 2007). Knockout mouse models of this disease have shown that absence of spatacsin leads to motor and cognitive impairments, consistent with symptoms observed in SPG11 patients (Branchu et al., 2017; Varga et al., 2015). Both models also revealed that loss of spatacsin promotes the accumulation of autophagic material in lysosomes, that was shown to result from the progressive accumulation of lipids, and notably simple gangliosides (Boutry et al., 2018).

Gangliosides are glycosphingolipids particularly enriched in the membrane of neurons (Xu et al., 2010). They can be internalized by endocytosis leading to their progressive degradation at the surface of intraluminal vesicles in lysosomes (Sandhoff and Sandhoff, 2018). Accumulation of gangliosides in lysosomes is observed in several lysosomal storage disorders (LSD)(Breiden and Sandhoff, 2019). Such accumulation can either be a direct consequence of mutations in genes encoding enzymes promoting gangliosides degradation, such as in Sandhoff or Tay Sachs diseases (Sandhoff and Harzer, 2013), or gangliosides may accumulate as a secondary consequence of other lysosomal metabolic impairment, as for example in Niemann Pick Type C disease (Breiden and Sandhoff, 2020). Most LSD that show accumulation of gangliosides in lysosomes also present with neurodegeneration (Sandhoff and Harzer, 2013). Substrate reduction therapy has been proposed for some LSD to prevent accumulation of gangliosides in lysosomes and delay disease progression (Jeyakumar et al., 1999; Marshall et al., 2019; Platt et al., 1997). In SPG11, the cause of ganglioside accumulation is unknown. However, we demonstrated in an *in vitro* study that decreasing ganglioside synthesis reduced neuronal death in primary cultures of neurons derived from a *Spg11* knockout (*Spg11^-/-^*) mouse model and improved motor behavior in a *Spg11* zebrafish model (Boutry et al., 2018), supporting the potential of substrate reduction therapy in SPG11.

Gangliosides are derived from lactosylceramide, which is converted into the simplest ganglioside GM3 by an enzyme encoded by the gene *St3gal5* (Sandhoff and Sandhoff, 2018). All other gangliosides from the a, b and c series are derived from GM3. We thus hypothesized that downregulation of *St3gal5* could prevent the accumulation of gangliosides, and provide a proof of concept that substrate reduction therapy delays onset of symptoms in a mouse model of SPG11. We subsequently evaluated substrate reduction therapy using a pharmacological inhibitor of glucosylceramide synthase that can cross the blood-brain barrier, venglustat.

## Material and methods

### Ethical Approval

The care and treatment of animals were in accordance with European Union Directive 2010/63/EU and with national authority (Ministère de l’Agriculture, France) guidelines for the detention, use, and ethical treatment of laboratory animals. All the experiments were approved by the local ethics committee (APAFIS, reference #201604201549915 for Paris Brain Institute mouse cohort, and references #25408-202005111212761 v7 and #30243- 2021022616138562 v4 for Dynacure mouse cohort), and experiments were conducted by authorized personnel in specific pathogen–free animal facilities. Patient-derived materials were obtained through procedures approved by the ethics committee with the written, informed consent of the family (approval SPATAX n° DC-2008-236 - RBM029 09/02/2004).

### Animals

The *Spg11* knockout (*Spg11^-/-^*) mouse model was previously described (Branchu et al., 2017). All mice used in the study were issued from crossing of heterozygous (*Spg11^+/-^*) mice; control *Spg11^+/+^* and knockout *Spg11^-/-^* mice used for all experiments were littermates. All analyses were performed by an evaluator bling to the genotype. Cognitive and motor function impairment were evaluated using the Y-maze and rotarod tests, respectively, following the protocols previously described (Branchu et al., 2017). Two independent cohorts of mice originating from the same colony were used in the study. The cohort that underwent treatment with AAV-PHP.eB viral vector was housed in the animal facility of Paris Brain Institute. Natural history study of this cohort demonstrated early onset cognitive and motor deficits, as previously described (Branchu et al., 2017). The second cohort, treated with venglustat was housed at Chronobiotron animal facility in Strasbourg, France. Natural history study in this independent cohort confirmed the occurrence of cognitive and motor deficit in *Spg11^-/-^* mice, albeit with slight differences (Supplementary Figure 1).

### Human brain

The autopsic SPG11 case (Patient IT) was previously described (Denora et al., 2016). Immunohistological analysis of the patient brain was performed on paraffin sections of cerebellum as described (Denora et al., 2016).

### Reagents

Venglustat was purchased from MedChemExpress (#HY-16743). The immunostaining were performed using mouse monoclonal anti-phosphorylated neurofilament (clone SMI31, Covance), rat monoclonal anti-Lamp1 (clone 1D4B, used for mouse samples, Development Studies Hybridoma Bank, University of Iowa, USA, deposited by JT August), monoclonal anti-Lamp1 (clone H5G11, used for human samples, Santa Cruz Biotechnology), chicken anti-GFP (#ab13970, Abcam), mouse anti-GM2 (hybridoma supernatant produced in-house) (Dobrenis et al., 1992; Natoli et al., 1986), and mouse anti-acetylated-α-tubulin (clone 6-11B- 1, #ab2410, Abcam). Fluorescent secondary antibodies for immunostaining (coupled to Alexa-488, Alexa-555 or Alexa 647) were purchased from Thermofisher.

AAV-PHP.eB production

AAV-PHP.eB vector expressing GFP together with a control miRNA (5’- AAATGTACTGCGCGTGGAGACGTTTTGGCCACTGACTGACGTCTCCACGCAGTAC ATTT -3’) or a miRNA downregulating mouse *St3Gal5* (5’- ATAACAGAGCCATAGCCGTCTGTTTTGGCCACTGACTGACAGACGGCTGGCTCTG TTAT -3’, previously described (Boutry et al., 2018)) under the control of CMV promoter were produced using HEK293 cells as previously described (Chan et al., 2017). Titration of viral particles was measured by qPCR as previously described (Aurnhammer et al., 2012).

#### Mouse treatment

*St3gal5* knockdown: 3.10^12^ genome copies of AAV-PHP.eB viral particles were diluted in 40 µl of PBS and injected intravenously in the caudal vein of mice at the age of 3 weeks. The viral dose was chosen after titration experiments.

Venglustat treatment: venglustat was mixed with food as previously established (Marshall et al., 2016) at 0.06 g/kg in A04 diet (Safe). This formulation provided approximately 12_mg of venglustat/kg/day for a 20-g mouse eating 4_g of food per day.

#### Viral genome copy analysis

Mice were killed by CO_2_, and brains were rapidly extracted to dissect cortex and hippocampus. Tissues were lysed to extract genome DNA. 20 ng of DNA was analyzed by qPCR using primers specific for vector DNA (*GFP*) and genomic DNA (*ADCK3*) on a LightCycler 480 II (Roche). Viral genomes per cell were calculated by dividing total viral genomes (detected by GFP) by diploid copies of the *ADCK3* gene (Liguore et al., 2019). The following primer sets were used: GFP: forward 5′- GAACGGCATCAAGGTGAACT-3′, reverse 5′- GAACTCCAGCAGGACCATGT-3’; AGCK3: Forward 5’- CCACCTCTCCTATGGGCAGA-3’; reverse 5’- CCGGGCCTTTTCAATGTCT-3’.

#### Histological evaluation

Mice were injected with a lethal dose of Euthasol (140/mg/kg) and were subjected to intracardiac perfusion of 4% paraformaldehyde (PFA) in PBS. The brains were dissected and post-fixed by incubation for 24 h in 4% PFA and 24 h in 30% sucrose PBS. 20 µm slices were cut on a freezing microtome (Microm HM450, Thermo Scientific) and maintained in 0.02% sodium azide in PBS at 4°C. After 90 min incubation in blocking solution, sections were incubated with primary antibodies in 2% BSA/0.25% Triton X-100 in PBS overnight at 4°C. After washing, the sections were incubated with the secondary antibodies for 90 min at room temperature, and mounted in Fluoromount-G mounting medium (Southern Biotechnology). Staining specificity was determined by incubation in the absence of primary antibodies. Analysis of the samples was performed in a blind manner.

#### *In situ* hybridization

*In situ* hybridization was performed using an RNAscope® Multiplex Fluorescent Reagent Kit v2 (Ref. 323100; Advanced Cell Diagnostics) according to the manufacturer’s instructions. The RNA probes (Mm-St3gal5, Ref. 821421; GFP-C2, Ref. 400281) were used to target mRNAs. In each experiment, we used Mm-Ppib (mouse peptidylprolyl isomerase B) and Dapb (bacterial dihydrodipicolinate reductase) probes as positive and negative controls, respectively. Opal fluorophores 520 and 650 (respectively Ref. PN FP1487001KT and PN FP1496001KT, Akoya Biosciences) were used to reveal mRNAs targets (*St3gal5* and *GFP* respectively). Fresh-frozen samples were sliced with a cryostat (Leica CM3050S) at 10 µm thickness. First, slices were fixed in PFA 4% solution for 30 min at 4°C, then rinsed in cold PBS during 10 min at 4°C twice. Samples were incubated with hydrogen peroxide for 10 min at room temperature. Slices were incubated with protease IV for 30 min at room temperature. Hybridization of the probes to the RNA targets was performed by incubation in the HybEZ Oven for 2 hr at 40°C. Then the slices were processed for standard signal amplification steps and fluorophore staining steps according to manufacturer’s instructions. Images were acquired with an inverted Leica SP8 confocal microscope (63X objective, N.A. 1.4).

Quantification was performed as previously described (Jolly et al., 2019). Probe signals were assigned to each nucleus by proximity, within a maximum cell radius of 10 µm. For each cell positive with GFP probe, we quantified the number of St3gal5 probes. A cell was considered positive for a staining when presenting at least one count. Cells positive for GFP were then classified in 5 classes according to the signal count for St3gal5 probes: cells negative for St3gal5, and 4 classes corresponding each quartile from minimum expression (0- 25%) to maximum expression (75-100%) for each experiment. These values were used to calculate an H-Score as follows: H-Score= (0×% negative cells)+(1×% of cells in 1^st^ quartile)+(2×% of cells in 2^nd^ quartile)+(3×% of cells in 3^rd^ quartile)+(4×% of cells in 4^th^ quartile), following recommendations by Cell Diagnostics USA (Newark, CA, USA) for the evaluation of RNAscope assays.

### Lipidomics

Lipidomic analysis was performed on the cortices of 4 month-old mice. Preparation of cortices, extraction of lipids and lipid analysis by liquid chromatography coupled with tandem mass spectrometry (LC-HRMS/MS) was performed as previously described (Boutry et al., 2018). The relative amount of each lipid was quantified as the area of its chromatographic peak, and it was normalized to the exact fresh weight of each cortex.

#### Quantification of neurofilament light chain (NfL) by Single Molecule Array (Simoa)

To quantify NfL in the mouse model, blood was sampled from the submandibular vein on mice anesthetized with isofluorane. After coagulation for 1 hour at 22°C, blood was centrifuged at 2,500 g for 30 min (22°C), and the serum was collected and stored at -80°C until analysis. Serum NfL levels were measured using the Simoa NF-Light Advantage Kit from Quanterix on the Simoa HD-1 Analyzer (Quanterix). Samples were diluted 1:20 in assay diluent and evaluated in duplicate following the manufacturer’s instructions.

To quantify NfL in SPG11 patients or healthy subject, we collected plasma samples on an EDTA tube anticoagulant from healthy controls and 21 patients presenting biallelic mutations in the *SPG11* gene. Plasma was frozen at -80°C, and stored in the local biobank. Plasma NfL levels were measured in duplicate using the aforementioned ultrasensitive Simoa assay, as previously established (Kuhle et al., 2019).

#### Primary culture of mouse neurons

Primary cultures of mouse cortical neurons were performed as previously described on E14.5 embryos obtained from crossing of heterozygous *Spg11^+/-^* mice (Pierga et al., 2023). Neurons were seeded at a density of 150,000 cells/cm^2^ on glass coverslips in 24-well plates and grown in Neurobasal medium supplemented with 2% B27 (ThermoFisher). Neurons were transfected with plasmids after 4 days of culture. For each well of 2cm^2^, we prepared a mix containing 1.2µl of Lipofectamine 2000 (ThermoFisher) with 1µg of Plasmid DNA in 30µl of Opti-MEM medium (ThermoFisher). The mix was added to cultured neurons for 3 hours, and neurons were analyzed after 8 days of culture.

#### Patient Fibroblasts and generation of human cortical neurons

Skin biopsies were collected from two healthy female subjects and two SPG11 female patients. Patient SPG11-1 (FSP-1123-004) carried two heterozygous truncating mutations *in trans* (c.2431 C>T, p. Gln811X; deletion of exon 29) and was previously described (Boutry et al., 2018). Patient SPG11-2 (FSP-446-014) carried a homozygous pathogenic variant (c.6100 C>T, p.R2034X). She had delayed acquisition of writing and reading and followed special schooling At age 10 there were no pyramidal signs. Gait difficulties started at age 17 and examination at age 19 showed increased reflexes in all limbs, bilateral extensor plantar reflex, as well as mild weakness distal in hands and feet. There were no cerebellar signs and ocular movements were normal. Nerve conduction was normal.

Fibroblasts were reprogrammed into iPS cells, and characterization of these cell lines were performed as previously described (Boutry et al., 2018). iPS cells were differentiated into forebrain neural progenitors and then into an enriched population of cortical neurons as previously described (Vazin et al., 2014) for 3 weeks. Venglustat was added at 5µM in the differentiation medium, and the medium was changed every 3-4 days.

#### Generation of isogenic iPS cell lines

Isogenic iPS cell lines were generated by introducing a homozygous stop mutation (c.6100 C>T, p.R2034X) in a line of iPS cells derived from an healthy subject (Boutry et al., 2018). gRNA and homology DNA repair (HDR) template were designed using the CRISPOR tool (Haeussler et al., 2016) and Benchling tool (https://benchling.com). The sequence of the selected gRNA was TGACCGATGCAAAcGAGCCC AGG, and the sequence of the single strand DNA oligonucleotide (ssODN) for HDR was AAGCCATGCTCCGGAAAATCTTGGCCTCTCAGCAGCCTGACCGATGCAAAtGAGCtCAGGCCTTCATCAGCACACAGGGCCTTAAGCCAGATACTGTGGC (both obtained from IDT). 10^6^ iPS cells were transfected using Nucleofector 4D and P3 Primary Cell 4D X transfection Kit (Amaxa) with the RNP complex composed of 225 pmol of each RNA (crRNA and tracrRNA-ATTO550) and 120 pmol of HiFiCas9 protein containing 3 NLS as well as 500 pmol ssODN as HDR template (all from IDT). 24 hours later, ATTO550 positive iPS cells were FACS sorted and plated at 10cells/cm^2^ on Laminin521 (StemCell) coated plates with CloneR supplement (StemCell) for clonal selection. After 7 days in culture, iPS cell clones were picked under a stereomicroscope and cultured in Laminin521-coated 96 well plates. When confluent, iPS cell clones were duplicated for cryopreservation and DNA extraction. Clones were then analyzed by PCR and Sanger sequencing. Quality control of positive clones and analysis of chromosome integrity were performed with the iCS-digital^TM^ Pluri solution (StemGenomics).

### Immunostaining

Cultured neurons were fixed in 4% PFA in PBS for 20 min and then permeabilized for 5 min in PBS containing 0.2% Triton X-100. Cells were then blocked for 30 min in PBS with 5% w/v BSA (PBS-BSA) and incubated with primary antibodies in PBS-BSA overnight at 4°C. Cells were washed three times with PBS and incubated with secondary antibodies coupled to fluorophores. After three washes with PBS, glass coverslips were then mounted on glass slides using Prolong Gold antifade reagent (Thermofisher) and imaged by confocal microscopy. Staining specificity was determined by incubation in the absence of primary antibodies.

### Imaging

Whole cortex, cerebellum or hippocampus images were obtained with a NanoZoomer 2.0-RS (Hamamatsu) equipped with a 20 × objective, N.A. 0.75. Identical brightness, contrast and colour balance adjustments were applied to all groups. Confocal images were acquired with an inverted Leica SP8 confocal laser scanning microscope, with a 63 × objective, N.A. 1.4. Quantification of cell populations and number of swellings were performed manually using ImageJ. For quantification of GM2 ganglioside level, regions of interest (ROIs) corresponding to the cell body of individual neurons were surrounded using ImageJ, and the mean of fluorescence intensity of ganglioside immunostaining in each ROI was quantified using the measure tool (Boutry et al., 2018). Values were normalized so that samples corresponding to *Spg11^+/+^* mice treated with control virus had a value of 1 in each experimental group.

### Statistics

All statistical tests were performed using GraphPad Prism 9 and the tests are described in the figure legends. Multiple comparisons with two different parameters (e.g. genotype and treatment) were performed using two-way ANOVAs.

## Results

Downregulation of *St3gal5* prevents accumulation of gangliosides in *Spg11^-/-^* mice We used a miRNA approach to downregulate expression of *St3gal5*. A miRNA targeting *St3gal5*, or a control sequence, was introduced into the 3’ untranslated region of the GFP mRNA (Figure 1A), allowing the detection of cells expressing the miRNA (Boutry et al., 2018). Both constructs were used to generate AAV-PHP.eB viral vectors that allow expression of the transgene under the control of the CMV promoter in the central nervous system upon intravenous injection (Chan et al., 2017). We injected 3.10^12^ genome copies of AAV-PHP.eB particles expressing either the control or the *St3gal5* miRNA (Figure 1B) at the age of 3 weeks in control (*Spg11^+/+^*) or *Spg11* knockout (*Spg11^-/-^*) mice, before onset of symptoms (Branchu et al., 2017). We first quantified at the age of 6 weeks the number of AAV-PHP.eB viral genome copies (VGC) per cell in the cortex and hippocampus using qPCR to evaluated the efficiency of gene transfer (Liguore et al., 2019). This analysis showed that all groups of mice incorporated similar number of viral genomes in the cortex and the hippocampus (Supplementary Figure 2A,B). We then evaluated the expression of the transgene by quantifying the proportion of neurons expressing GFP in various brain regions affected by the disease (Branchu et al., 2017). Purkinje cells in the cerebellum and granule cells of CA3 region in the hippocampus were efficiently transduced upon injection of the viral vectors in the blood circulation (Supplementary Figure 2C,D). The proportion of Purkinje cells expressing GFP however dropped by the age of 8 months. Cortical neurons also expressed GFP, although the proportion of transduced neurons was lower than in the cerebellum or hippocampus (Supplementary Figure 2E).

**Figure 1.**
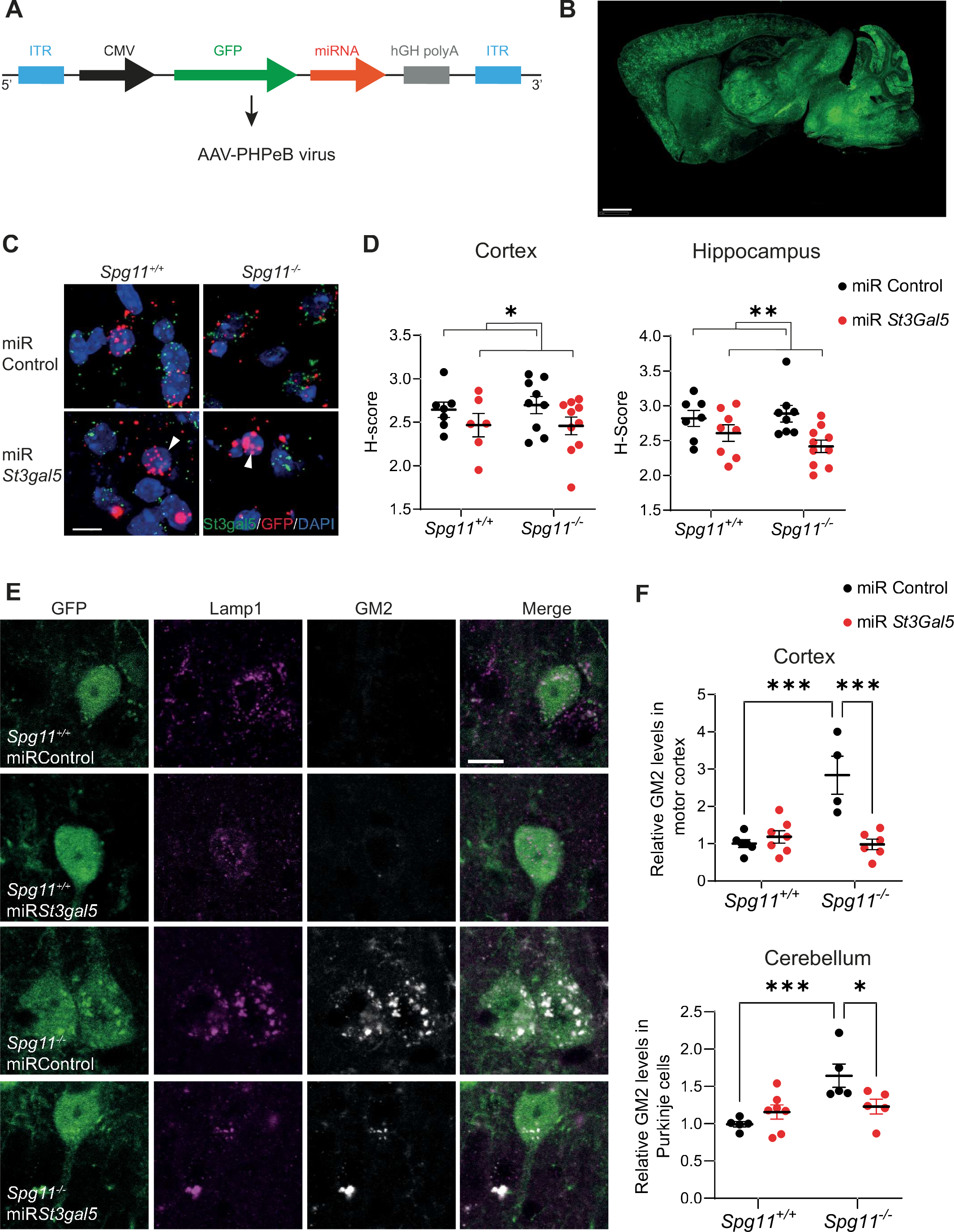
Downregulation of *St3gal5* delays accumulation of gangliosides in neurons of *Spg11^-/-^* mice. (**A**) Scheme illustrating the constructs used to generate AAV-PHP.eB virus. The sequences of the control miRNA or the miRNA downregulating *St3gal5* were inserted in the 3’ untranslated region of the *GFP* mRNA. The construct is under the control of cytomegalovirus (CMV) promoter to allow constitutive and high expression. Human growth hormone (hGH) polyadenylation signal was used for termination and polyadenylation of the transcript. The sequences were flanked with inverted terminal repeat (ITR) sequences allowing its insertion into AAV-PHP.eB viral particles. (**B**) Sagittal section of the brain of a mouse 3 weeks after intravenous injection of 3.10^12^ genome copies of AAV-PHP.eB expressing GFP, showing the widespread expression of the transgene. Scale bar 1mm. (**C**) Confocal images of RNA Scope staining in CA1 hippocampal neurons of *Spg11^+/+^* and *Spg11^-/-^* mouse injected with a control virus or AAV-PHP.eB downregulating *St3gal5* (miR *St3gal5*). Note the decrease in the number of *St3gal5* mRNA puncta (green) in cells expressing the miRNA downregulating *St3gal5* together with GFP (red puncta). Scale bar 5 µm. (**D**) Quantification of number of *St3gal5* dots (H-Score) in neurons positive for GFP in cortex and hippocampus. Means and SEM. N=6-10 independent mice. *P<0.05; **P<0.01; Two-way ANOVA. (**E**) Confocal images of GFP, Lamp1 and GM2 ganglioside in neurons of motor cortex of *Spg11^+/+^*and *Spg11^-/-^* mouse injected with a control virus or AAV-PHP.eB downregulating *St3gal5*. Scale bar 10µm. (**F**) Quantification of the relative levels of GM2 ganglioside in neurons of cortex (Top) or cerebellum of 4 month-old *Spg11^+/+^* and *Spg11^-/-^*mouse injected with a control virus or AAV-PHP.eB downregulating *St3gal5*. Means and SEM. N= 4-7 independent mice/group. *P<0.05; **P<0.01; *** P<0.001; Two-way ANOVA followed by Holm-Sidak multiple comparison test.

We then evaluated the *in vivo* efficiency of the miRNA downregulating *St3gal5* by a quantitative *in situ* hybridization approach. This method allowed to quantify expression levels of *St3gal5* in transduced neurons that were positive for GFP (Figure 1C). It showed that the expression of the miRNA slightly and significantly reduced the expression of *St3gal5* in transduced cortical and hippocampal neurons compared to the control virus (Figure 1D). Previous data showed that knocking out *Spg11* led to accumulation of simple gangliosides in neurons, notably GM2 (Boutry et al., 2018). Downregulation of *St3gal5* reduced the intensity of the immunostaining of the ganglioside GM2 in neurons of the cortex, cerebellum and hippocampus of *Spg11^-/-^*mice at the age of 4 months (Figure 1E-F; Supplementary Figure 3A). However, by the age of 8 months, downregulation of *St3gal5* only marginally reduced levels of GM2 immunostaining in the cortex of *Spg11^-/-^* mice (Supplementary Figure 3B).

Together, these data demonstrate that injection of an AAV-PHP.eB vector expressing a miRNA targeting *St3gal5* delayed the accumulation of gangliosides in neurons in the brain of *Spg11^-/-^* mice.

Downregulation of *St3gal5* delays motor and cognitive symptom onset in *Spg11^-/-^* mice Next, we explored whether downregulation of *St3gal5* may prevent or delay the onset of cognitive or motor symptoms observed in *Spg11^-/-^* mice (Branchu et al., 2017) using the Y-maze spontaneous-alternation test and the rotarod test, respectively. These tests previously showed significant decrease in the performance of *Spg11^-/-^* mice compared to *Spg11^+/+^* mice at the age of 4 and 8 months (Branchu et al., 2017).

Downregulation of *St3gal5* did not change the performance of *Spg11^+/+^* mice in Y-maze and rotarod tests compared to mice injected with the control virus (Figure 2A and 2B). *Spg11^-/-^* mice injected with the AAV-PHP.eB expressing the miRNA downregulating *St3gal5* had a higher performance than *Spg11^-/-^*mice injected with the control virus at the age of 6 weeks and 4 months in the Y-maze test, and at the age of 4 months in the rotarod test (Figure 2A and 2B). However, downregulation of *St3gal5* did not provide any significant improvement at the age of 8 months (Figure 2A and 2B), corresponding to reduced transgene expression observed at this age (Supplementary Figure 2D). Overall, these results demonstrated downregulation of *St3gal5* delayed the onset of cognitive and motor symptoms in *Spg11^-/-^* mice.

**Figure 2.**
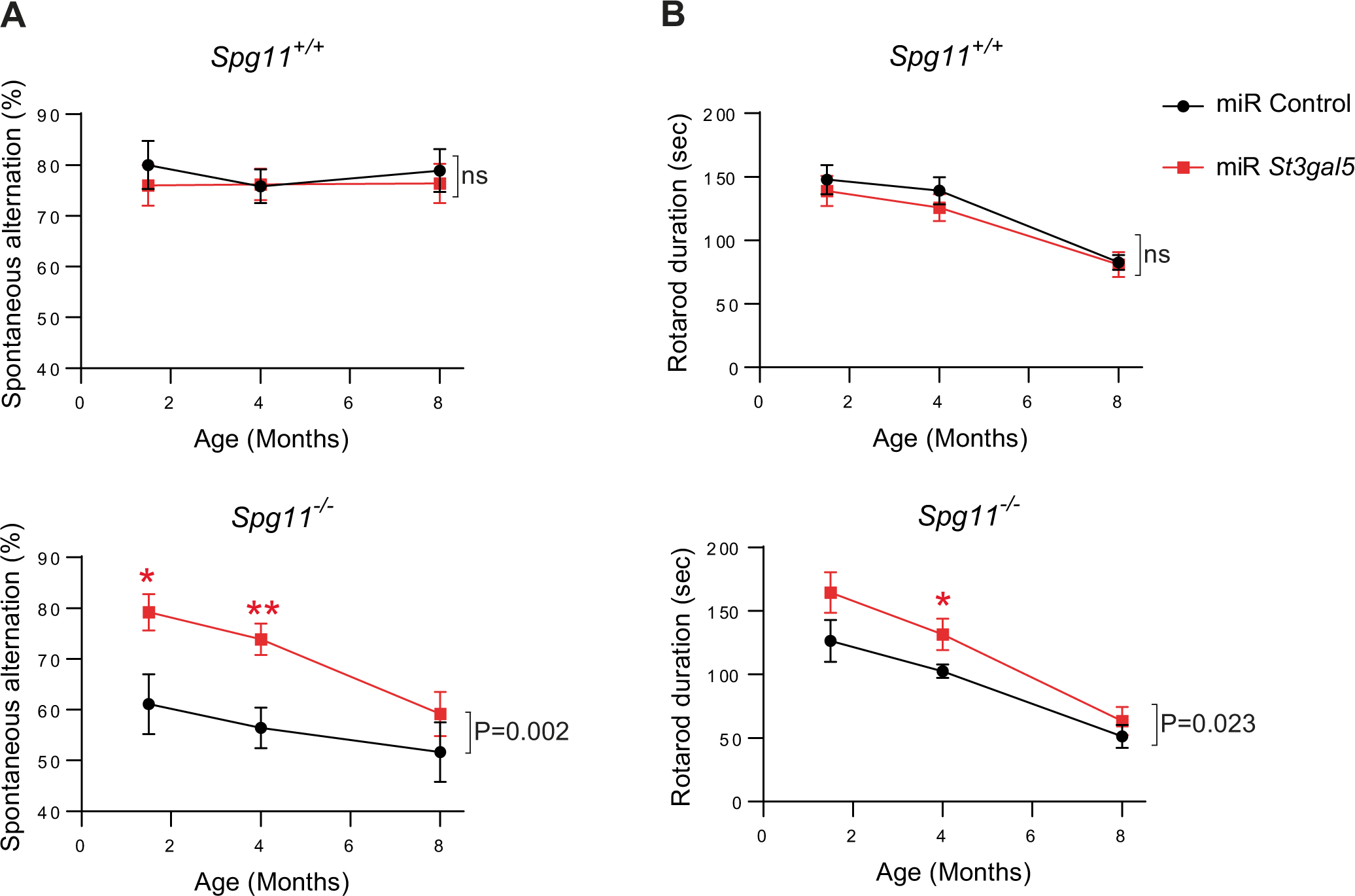
Downregulation of *St3gal5* delays motor and cognitive symptom onset in *Spg11^-/-^* mice. Evaluation of the cognitive function using the Y-maze test (**A**) and the motor function using the accelerating rotarod test (**B**) in *Spg11^+/+^* (Top) and *Spg11^-/-^* mice (Bottom) injected with a control virus or with AAV-PHP.eB downregulating *St3gal5*. Means and SEM, N>9 mice/group. *P<0.05; **P<0.01; Two-way ANOVA followed by Holm-Sidak multiple comparison test. P values shown on the graphs indicate statistical differences between the groups of mice injected with control or AAV-PHP.eB downregulating *St3gal5* (treatment effect).

Downregulation of *St3gal5* restores abnormal axonal trafficking of lysosomes in *Spg11^-/-^*

### neurons

The absence of the *Spg11* product, spatacsin, was recently shown to impair the axonal trafficking of lysosomes *in vitro* (Pierga et al., 2023). Impaired axonal trafficking has previously been implicated in many motor neuron diseases, including HSP (De Vos et al., 2008). Such trafficking defects were revealed by the presence of axonal swellings or spheroids in models of SPG4, SPG5 or SPG7 (Ferreirinha et al., 2004; Mou et al., 2023; Tarrade et al., 2006). Transfection of primary cultures of cortical neurons with a plasmid expressing GFP revealed higher number of swellings in neurites of *Spg11^-/-^* compared to *Spg11^+/+^* neurons (Figure 3A). Immunostaining demonstrated that these swellings in *Spg11^-/-^* neurons were positive for the late endosome or lysosome marker Lamp1 as well as for the ganglioside GM2 (Figure 3B), suggesting that they are a consequence of impaired degradation of gangliosides by lysosomes. To test this hypothesis, we investigated whether downregulation of *St3gal5* may compensate this cellular dysfunction. Transfection of *Spg11^-/-^*primary neurons with a plasmid expressing GFP and the miRNA targeting *St3Gal5* decreased the number of swellings in *Spg11^-/-^*neurons (Figure 3C). To confirm the link between gangliosides and the number of swellings, we inhibited ganglioside degradation by downregulating *Neu1*, which encodes a neuraminidase and promotes accumulation of gangliosides in lysosomes (Boutry et al., 2018). Downregulation of *Neu1* led to a higher proportion of neurons presenting swellings in primary cultures (Supplementary Figure 4A).

**Figure 3.**
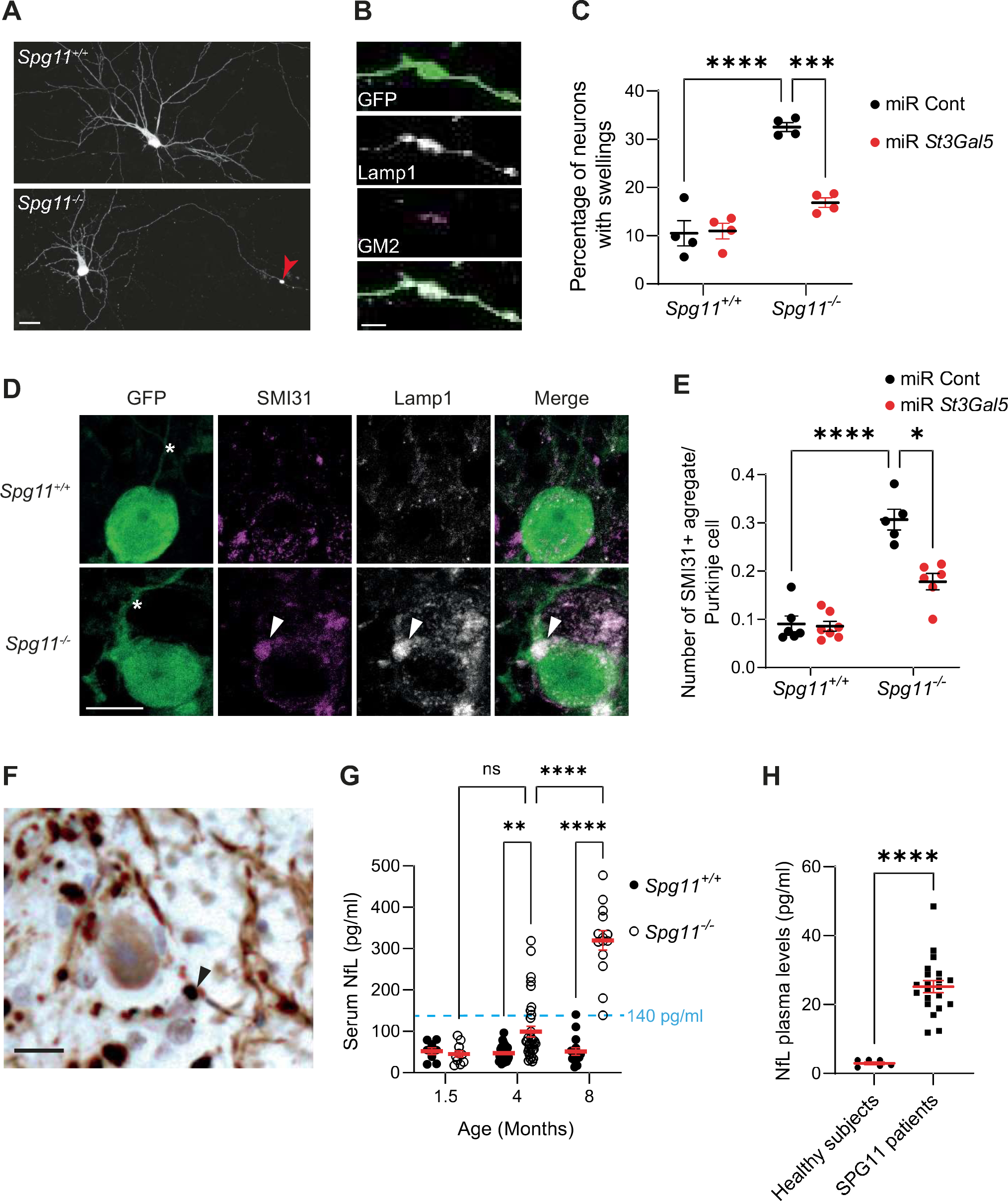
Downregulation of *St3gal5* prevents swellings in *Spg11^-/-^* neurons. (**A**) Confocal images of primary cultures of *Spg11^+/+^* and *Spg11^-/-^* mouse neurons transfected with a vector expressing GFP. Note the presence of a swelling in neurites of *Spg11^-/-^*neuron (red arrowhead). Scale bar 20 µm. (**B**) High resolution confocal image of a swelling in a *Spg11^-/-^*neuron identified by the presence of GFP and immunostained for GM2 and the lysosomal marker Lamp1. Scale bar 5µm. (**C**) Quantification of the proportion of *Spg11^+/+^* or *Spg11^-/-^* neurons presenting swellings when transfected with vectors expressing GFP and either a control miRNA or miRNA downregulating *St3Gal5*. Means and SEM. N= 4 independent experiments with more than 50 neurons analyzed per experiment. *** P<0.001; **** P<0.0001; Two-way ANOVA followed by Holm-Sidak multiple comparison test. (**D**) Cerebellum of 4 month-old *Spg11^+/+^* and *Spg11^-/-^* mice injected with control AAV-PHP.eB immunostained with GFP, Lamp1 and SMI31 antibodies. Axons of Purkinje cells (asterisks) were identified by their orientation toward the granule layer of the cerebellum. Note the presence of axonal swelling positive for Lamp1 and SMI31 in Purkinje cell of *Spg11^-/-^* mouse (white arrowheads). Scale bar: 10 µm. (**E**) Quantification of the proportion of Purkinje cells presenting axonal swellings in 4 month-old *Spg11^+/+^* and *Spg11^-/-^* mice injected with control or AAV-PHP.eB downregulating *St3gal5*. Means and SEM. N=5-7 independent mice. *P<0.05; **** P<0.0001. Two-way ANOVA followed by Holm-Sidak multiple comparison test. (**F**) SMI31 immunostaining of cerebellum slice of the autopsy brain of a SPG11 patient. Note the presence of axonal swelling labelled by phosphorylated neurofilaments (black arrowhead). Scale bar 10µm. (**G**) Quantification of the serum levels of NfL in *Spg11^+/+^* and *Spg11^-/-^*mice at the age of 1.5, 4 and 8 months. Note that by the age of 8 months all *Spg11^-/-^*mice except one present NfL levels higher than 140 pg/ml, whereas *Spg11^+/+^*mice (except one) had values below this threshold. Means and SEM. N> 10 mice per group. **P<0.01; ****P<0.0001. Two way ANOVA followed by Holm-Sidak multiple comparison test. (**H**) Quantification of NfL levels in plasma of SPG11 patients and aged-matched healthy subjects. **** P<0.0001. Mann-Whitney test.

We then investigated whether neurite swellings also occurred in the brains of symptomatic *Spg11^-/-^* mice. We focused on Purkinje cells, which are amongst the most affected in *Spg11^-/-^* mice (Branchu et al., 2017). Consistent with previous results, we observed no neuronal death of Purkinje cells in *Spg11^-/-^* mice by the age of 4 months despite the presence of motor and cognitive symptoms (Supplementary Figure 4B). However, we observed an accumulation of Lamp1 immunostaining in the proximal part of the axons of Purkinje cells of 4-month-old *Spg11^-/-^* mice (Figure 3D), which is consistent with impaired axonal transport. These structures were positive for the GM2 ganglioside (Supplementary Figure 4C) as well as for the phosphorylated neurofilament heavy chain marker SMI31 (Figure 3D) labelling axonal spheroids that have been associated with impaired axonal trafficking (Coleman, 2005; Sharma et al., 2020). The proportion of Purkinje cells with axonal accumulation of SMI31 was lower at the age of 4 months when *Spg11^-/-^*mice were injected with AAV-PHP.eB downregulating *St3gal5* (Figure 3E), showing that preventing the accumulation of ganglioside reduced the number of axonal spheroids. Interestingly, analysis of the cerebellum of a SPG11 patient previously described (Denora et al., 2016) revealed dystrophic axons labelled by SMI31 (Figure 3F). Together these data suggest that loss of function mutations in SPG11 lead to formation of axonal spheroids, previously associated with impaired trafficking (Sharma et al., 2020), which is partially prevented by decreasing ganglioside synthesis.

### Downregulation of *St3gal5* lowers serum levels of neurofilament light chain in *Spg11^-/-^* mice

Quantification of neurofilament light (NfL) chain has been used as a biomarker in multiple neurodegenerative diseases (Coarelli et al., 2021; Khalil et al., 2018; Parnetti et al., 2019; Welford et al., 2022). We first evaluated NfL levels in the serum of *Spg11^+/+^* and *Spg11^-/-^* mice at 6 weeks, 4 months and 8 months. We noticed that the average levels of NfL progressively increased in serum of *Spg11^-/-^* mice (Figure 3G). At the age of 8 months, when *Spg11^-/-^* mice present overt neuronal death (Branchu et al., 2017), there was a clear discrimination between NfL levels in the serum of *Spg11^+/+^* and *Spg11^-/-^* mice, with a threshold of 140 pg/ml allowing to separate both groups. We therefore considered NfL levels higher than 140 pg/ml as a marker of the disease. Strikingly, even at the age of 4 months, some *Spg11^-/-^* mice presented levels of serum NfL higher than 140 pg/ml, suggesting a high variability in the disease progression amongst *Spg11^-/-^* mice (Figure 3G). Interestingly, NfL levels were also significantly higher in plasmas from symptomatic SPG11 patients than in healthy subjects (Figure 3H), suggesting that NfL may be used as a cross-species biomarker.

We then evaluated whether downregulation of *St3gal5* had an impact on serum NfL levels. Downregulation of *St3gal5* had no impact on the serum NfL levels in *Spg11^+/+^* mice (Table 1). In contrast, in *Spg11^-/-^*mice, downregulation of *St3gal5* significantly reduced the proportion of animals with NfL levels higher than the threshold of 140 pg/ml (Table 1), suggesting that downregulation of ganglioside synthesis had a measurable impact in the serum of mice and could be used as a measure for response to treatment.

**Table 1.** Proportion of mice with serum NfL>140 pg/ml at the age of 4.5 months.

| | miR Control | miR *St3gal5* | P value ($\chi^2$ test) |
| --- | --- | --- | --- |
| *Spg11*<sup>+/+</sup> | 0/16 | 0/15 | NS |
| *Spg11*<sup>-/-</sup> | 8/19 | 2/18 | 0.034 |

### Therapeutic action of venglustat

Downregulation of *St3gal5* demonstrated that decreasing ganglioside synthesis may be a therapeutic option to delay the onset of symptoms in SPG11. Pharmacological agents acting on glucosylceramide synthase, upstream of St3gal5, have been shown to prevent ganglioside accumulation in several mouse models of gangliosidoses (Jeyakumar et al., 1999; Marshall et al., 2019; Platt et al., 1997). Recently, the development of such agents that are crossing the blood brain barrier opened new avenues for therapy of some lysosomal storage disorders. We therefore investigated whether such an agent, venglustat, could be a therapeutic option for SPG11.

*Spg11^-/-^* mice were treated with venglustat (12mg/kg/day, see methods) from the age of 7 weeks and were evaluated at the age of 6 weeks (prior treatment) as well as 11 and 15 weeks for cognitive (Y-Maze) and motor (Rotarod) performance. At the age of 16 weeks, serum of mice was collected to evaluate levels of NfL as biomarkers. Half of the mice were perfused to perform histological investigations in the brain, and the other half of mice were used to collect tissues for lipidomic analysis. Venglustat inhibits the glucosylceramide synthase, preventing the conversion of ceramide into glucosylceramide (Figure 4A). Lipidomic analysis showed that venglustat treatment slightly but significantly lowered the levels of glycosphingolipids that are produced downstream of glucosylceramide synthase (Figure 4B).

**Figure 4.**
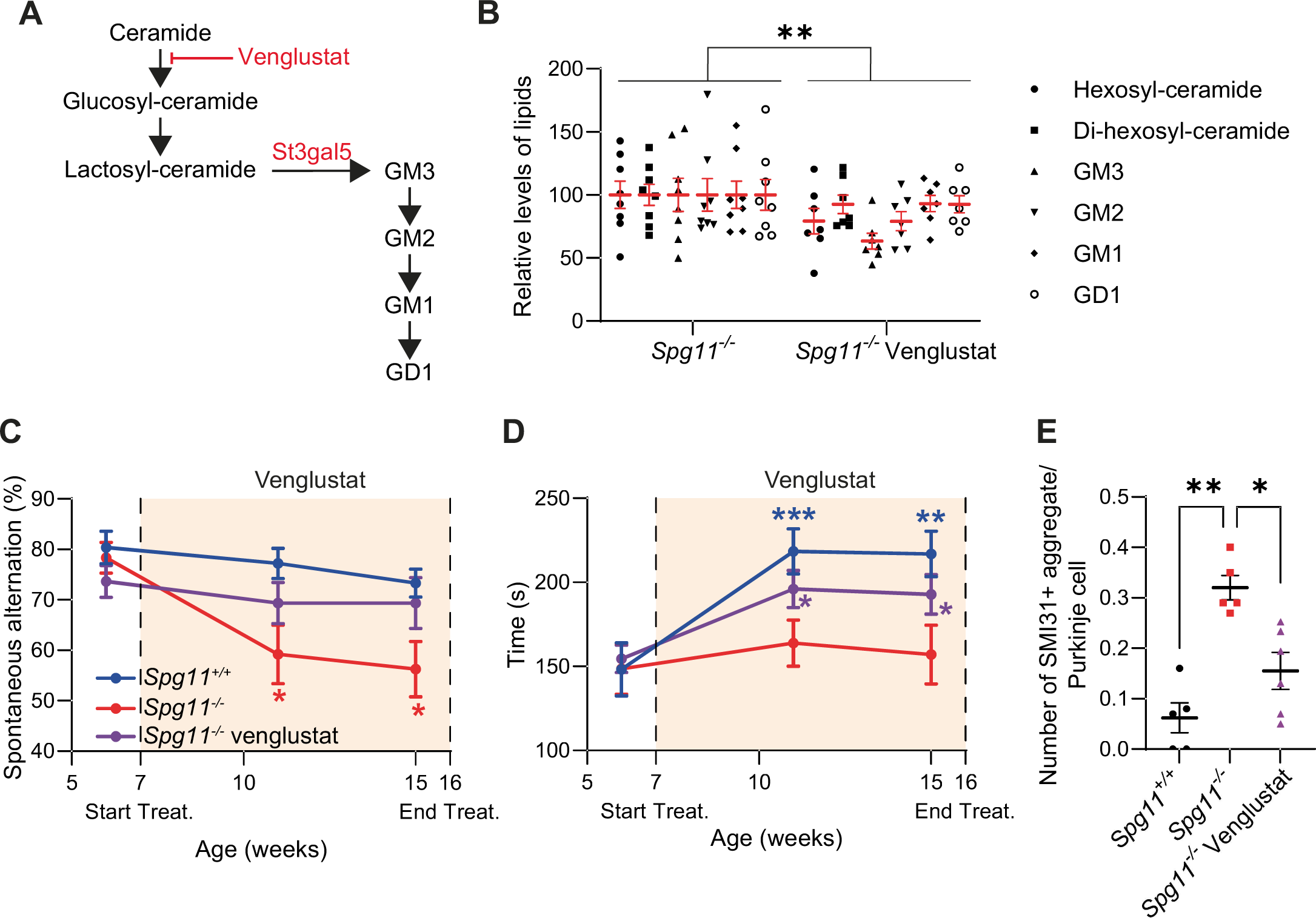
Treatment of *Spg11^-/-^*mice with venglustat delays disease progression. (**A**) Scheme showing the biosynthetic pathway of gangliosides (GM3, GM2, GM1, GD1) from ceramide. Venglustat is an inhibitor of glucosylceramide synthase converting ceramide into glucosyl-ceramide. (**B**) Quantification of lipids produced downstream of glucosylceramide synthase in the cortex of *Spg11^-/-^* mice and *Spg11^-/-^* mice treated with venglustat by lipidomic analysis. Mean and SEM. N>7 cortices/groups. **P<0.01, Two Way ANOVA. Note that glucosyl-ceramide and lactosyl ceramide were monitored within hexosyl-ceramide and di- hexosyl-ceramide. (**C, D**) Evaluation of the cognitive function by the Y-maze test (**C**) and the motor function by the accelerating rotarod test (**D**) in *Spg11^+/+^*mice, *Spg11^-/-^* mice and *Spg11^-/-^* mice treated with venglustat from the age of 7 weeks (colored zone). Data represent Mean and SEM at each time point. N> 15 mice/group. *P<0.05; **P<0.01; ***P<0.001 when compared to 6-week-old animals of the same group. Two Way ANOVA with repeated measures, followed by Holm-Sidak multiple comparison test. (**E**) Quantification of the proportion of Purkinje cells presenting axonal swellings in *Spg11^+/+^*, *Spg11^-/-^* mice and in *Spg11^-/-^* mice treated with venglustat. Means and SEM. N>5 mice. *P<0.05; **P<0.01. Kruskall Wallis test followed by Dunn’s multiple comparison test.

This cohort of mice, evaluated in a different facility with independent experimenters, confirmed the occurrence of significant cognitive and motor deficit in *Spg11^-/-^* mice, albeit with slight differences (Supplementary Figure 1). The cognitive performance of *Spg11^-/-^* mice significantly decreased with aging, whereas the performance of wild-type mice was stable. Treatment of *Spg11^-/-^* mice with venglustat prevented the decrease in cognitive performance (Figure 4C). The motor performance significantly increased with age in the *Spg11^+/+^* but not in the *Spg11^-/-^* mice. Venglustat treatment allowed a slight but significant increase of motor performance in *Spg11^-/-^* mice at the age of 11 and 15 weeks (Figure 4D). These data therefore suggested that treatment of *Spg11^-/-^* mice at an early stage of the disease delays onset of motor and cognitive signs.

Analysis of NfL in the serum of mice at the age of 16 weeks showed that treatment of *Spg11^-/-^* mice with venglustat prevented serum NfL to reach levels higher than 140 pg/ml (Table 2), similar to the observation made upon downregulation of *St3gal5*, although the difference between treated and untreated *Spg11^-/-^* mice was not significant. Finally, we evaluated whether venglustat may delay onset of the disease by restoring trafficking similarly to downregulation of *St3gal5*. Venglustat treatment in *Spg11^-/-^* mice led to a significantly lower proportion of Purkinje cells with axonal spheroids positive for SMI31 (Figure 4E), suggesting that venglustat and downregulation of *St3gal5* may act on the same pathway.

**Table 2.**
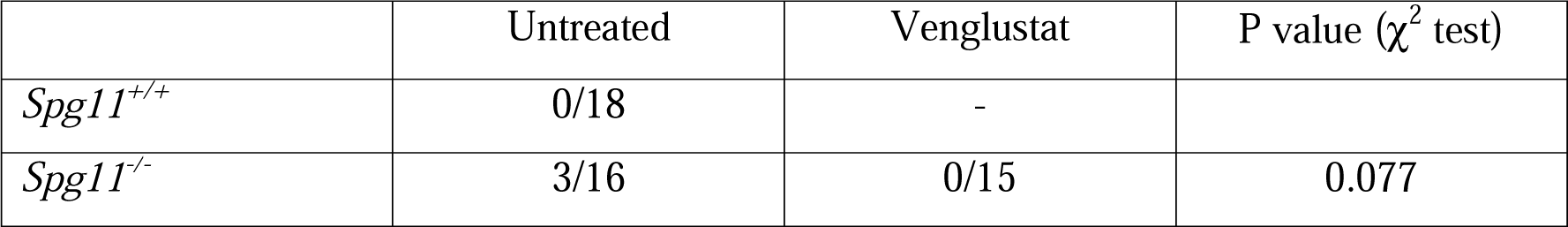
Proportion of mice with serum NfL>140 pg/ml at the age of 16 weeks.

| | Untreated | Venglustat | P value ($\chi^2$ test) |
| --- | --- | --- | --- |
| *Spg11*<sup>+/+</sup> | 0/18 | - |  |
| *Spg11*<sup>-/-</sup> | 3/16 | 0/15 | 0.077 |

Together, these data suggest that venglustat treatment (12mg/kg/day) delays the onset of motor and cognitive signs in *Spg11^-/-^* mice, which is reflected by a reduction in the number of Purkinje cells presenting axonal spheroids as well as in serum levels of NfL as biomarkers.

#### Venglustat decreases the number of neuronal swellings in human neurons

Finally, we evaluated whether venglustat may be effective on human neurons. Neurons with cortical identity were differentiated from induced pluripotent stem (iPS) cells obtained from two independent healthy subjects or SPG11 patients. Furthermore, we also used an isogenic iPS cell line in which a homozygous *SPG11* truncating mutation (c.6100 C>T, p.R2034X) was introduced by genome editing. After 3 weeks of differentiation, SPG11 neurons exhibited neurites swellings revealed by acetylated tubulin, similar to the observations in SPG4 neurons (Denton et al., 2014; Havlicek et al., 2014). Similar to the swellings observed in the neurons of *Spg11^-/-^* mice, swellings present in human SPG11 neurons were positive for the late endosomes or lysosomes marker Lamp1 as assessed by immunocytochemistry (Figure 5A). Treatment of human SPG11 neurons with venglustat during their differentiation significantly decreased the proportion of neurite swellings (Figure 5B-C), suggesting that venglustat is efficient on human SPG11 neurons as well.

**Figure 5.**
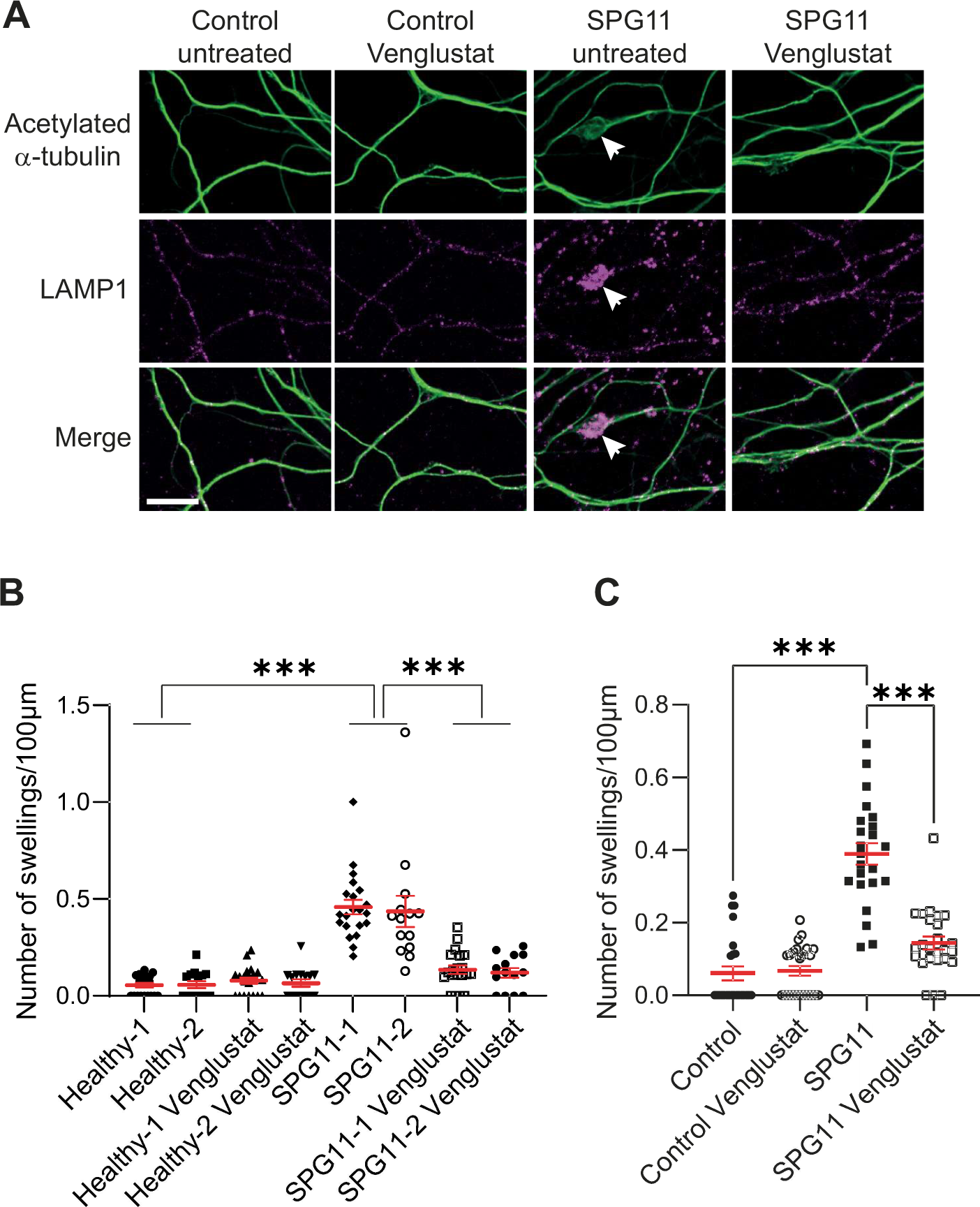
Venglustat prevents formation of neurite swellings in human neurons. (**A**) Acetylated tubulin and LAMP1 immunostaining of human neurons derived from iPS cells of a heathy subject (Control) and a SPG11 patient treated with a vehicle or with 5 µM venglustat for 3 weeks. The white arrows indicate neurite swellings. Scale bar 10 µm. (**B**) Quantification of the number of swellings in neurons differentiated from iPS cells of 2 independent healthy subjects and 2 independent SPG11 patients treated with 5 µM venglustat for 3 weeks. Mean and SEM. N>14 microscopy fields analyzed in at least two independent experiments. *** P<0.001. One way ANOVA followed by Holm-Sidak multiple comparison test. (**C**) Quantification of the number of swellings in neurons differentiated from a control iPS cell line or an isogenic line containing the same truncating *SPG11* mutation as patient SPG11-2 (c.6100 C>T, p.R2034X). Mean and SEM. N>23 microscopy fields analyzed in at least two independent experiments. *** P<0.001. One way ANOVA followed by Holm-Sidak multiple comparison test.

## Discussion

Hereditary spastic paraplegia type SPG11 is a neurodegenerative disease with currently no treatment option to prevent or delay disease progression. Here we show that decreasing ganglioside synthesis, either by downregulating the expression of *St3gal5* or by using a pharmacological inhibitor of glucosylceramide synthase, delays the onset of motor and cognitive signs in a mouse model of SPG11 and improves axonal damage in human and rodent neurons carrying SPG11 mutations.

Loss of function mutations in *Spg11* have previously been associated with accumulation of gangliosides (Boutry et al., 2018). However, this accumulation is modest (∼50% increase in global brain lipidomic analysis) compared to the accumulation observed in mouse models of gangliosidoses such as Sandhoff disease (several hundred folds increase (Marshall et al., 2019)). Substrate reduction therapies aiming at decreasing synthesis of gangliosides were proven to be efficient in mouse models of gangliosidoses (Jeyakumar et al., 1999; Marshall et al., 2019; Platt et al., 1997). The moderate accumulation of gangliosides in SPG11 suggested that mild reduction in synthesis of gangliosides may thus be sufficient to provide beneficial effects. As a proof of concept for our hypothesis, we used a miRNA to downregulate the expression of *St3gal5*. Complete loss of *ST3GAL5* in humans has been associated with severe neurodevelopmental disorders (Boccuto et al., 2014; Simpson et al., 2004), and we thus aimed for a moderate downregulation of this target. The RNA interference approach coupled to the use of AAV-PHP.eB viral vectors provided a modest but significant reduction in the expression of *St3gal5*, which was sufficient to delay the accumulation of GM2 ganglioside in lysosomes, as well as the onset of motor and cognitive symptoms. It was however not able to completely prevent the disease progression. The lack of long term protection may be explained by the minimal *St3gal5* downregulation that we achieved. Alternatively, it may be due to the progressive decrease in the expression of the transgene observed in some brain regions at the age of 8 months. Indeed, the transgene was expressed under the control of CMV promoter that has been shown to be progressively inactivated *in vivo* (Yew et al., 2001). Yet, the delay observed both in accumulation of gangliosides and in onset of motor and cognitive deficits supports the implication of ganglioside accumulation in disease progression.

The small molecule venglustat that inhibits an enzyme upstream of *St3gal5* also slowed down disease progression, suggesting that targeting the biosynthetic pathway of gangliosides is an effective therapeutic option for SPG11. Venglustat is an inhibitor of glucosylceramide synthase that crosses the blood brain barrier and that has proven to delay the onset of symptoms in mouse models of Sandhoff disease, neuronopathic Gaucher disease, as well as glucocerebrosidase (GBA)-related synucleinopathies (Marshall et al., 2019, 2016; Schidlitzki et al., 2023; Viel et al., 2021). Clinical data suggested safety of the molecule in healthy volunteers as well as Parkinson’s disease and Fabry disease patients (Deegan et al., 2023; Peterschmitt et al., 2022, 2021). Our preclinical study shows that a dose of 12 mg/kg/day of venglustat was sufficient to delay onset of symptoms in the mouse model of SPG11. The efficacy of a low dose of venglustat in our study is likely due to the relatively modest accumulation of gangliosides observed in the *Spg11^-/-^* model. The use of a low dose of venglustat may also avoid or decrease potential adverse effects that were observed in another study in mice (Schidlitzki et al., 2023): a dose of 60 mg/kg/day led to worsening of motor performance, suggesting possible adverse or off-target effects of the molecule (Schidlitzki et al., 2023). The behavioral benefit provided by venglustat treatment was weaker than the one provided by downregulation of *St3gal5*. This difference may be explained by a difference in the initiation of treatment. Treatment with venglustat was started from the age of 7 weeks whereas downregulation of *St3gal5* was initiated at the age of 3 weeks. This observation argues for initiating a treatment targeting ganglioside synthesis as early as possible.

The cohorts of mice treated with AAV.PHP.eB or venglustat presented slight differences in motor and cognitive deficits. This is likely due to the use of two independent groups of *Spg11^-/-^* mice in different facilities and evaluated by different experimenters (von Kortzfleisch et al., 2022). Despite these mild differences, the beneficial action of *St3gal5* downregulation or venglustat treatment was evident before the age of 4 months, ahead of the degeneration of neuron somas that was detected from the age of 8 months in the Spg11 mouse model (Branchu et al., 2017). This observation suggests that these therapeutic approaches may prevent neuronal dysfunction, long before neuronal death occurs. In a recent study, we identified that loss of function mutations in *Spg11* resulted in impaired trafficking of lysosomes in axons (Pierga et al., 2023), suggesting that alteration of intracellular trafficking may contribute to symptoms, as observed in many motor neuron diseases (De Vos et al., 2008). Pathogenic variants in genes encoding motor proteins such as kinesins KIF5A, KIF1A or KIF1C indicated that impaired axonal trafficking contribute to pathophysiology in several forms of HSP (Dor et al., 2014; Erlich et al., 2011; Reid et al., 2002). Furthermore, impaired axonal trafficking has also been observed in axons of *Spg4* knockout mice and in neurons derived from SPG4 iPS cells (Denton et al., 2014; Fassier et al., 2012; Havlicek et al., 2014), and they were associated with axonal swellings, suggesting that the latter could be a sign of impaired trafficking (Denton et al., 2014; Fassier et al., 2012). Similar swellings have also been observed in *Spg7* knockout mice and recently in human SPG5 neurons (Ferreirinha et al., 2004; Mou et al., 2023), suggesting that they may represent a commonality among HSP models. We reveal the presence of swellings in cultured human and mouse SPG11 neurons. Importantly, we also identified axonal swellings in Purkinje cells in *Spg11* knockout mice and in a SPG11 patient by the formation of phosphorylated neurofilaments spheroids that were previously associated with disrupted axonal transport in amyotrophic lateral sclerosis patients (Sharma et al., 2020). Decreasing ganglioside synthesis was sufficient to reduce the number of swellings in mouse and human neurons, suggesting that lysosomal dysfunction caused by ganglioside accumulation may alter lysosomal trafficking. Similarly, ameliorating lysosomal metabolic disturbances was also shown to restore trafficking of lysosomes in models of Niemann Pick type C or Sanfilippo disease (Beard et al., 2017; Roney et al., 2021). Therefore, the reduction in ganglioside biosynthesis in SPG11 may contribute to delay the onset of symptoms by restoring trafficking of lysosomes.

The levels of NfL that are used as a biomarker for many neurodegenerative diseases (Coarelli et al., 2021; Khalil et al., 2018; Parnetti et al., 2019; Welford et al., 2022) were higher in the plasma of symptomatic SPG11 patients compared to healthy subjects. They were also increased in the serum of *Spg11* knockout mice from the age of 4 months, before any loss of neuron somas is detected (Branchu et al., 2017). This suggests that peripheral NfL levels may reflect neuronal dysfunction that occurs before death of neuron somas, similar to an observation made in a mouse model of spinocerebellar ataxia SCA3 (Wilke et al., 2020). Importantly, downregulation of *St3gal5* that restored axonal trafficking, prevented NfL to reach high levels in the serum of *Spg11* knockout mice. Peripheral levels of NfL may thus be considered as a biomarker of the response to treatment. However, levels of NfL were highly variable in *Spg11* knockout mice, with some 6-week-old untreated knockout mice presenting NfL levels as high as 8-month-old untreated mice, despite a clear difference in the motor and cognitive performance. This observation calls for a thorough characterization of NfL levels over time, which may refine their use as a biomarker of response to treatment.

Together, our data show that decreasing ganglioside synthesis is a therapeutic option to delay disease onset in SPG11. The beneficial action of such treatment likely relies on restoring impaired axonal trafficking, including in human neurons. These results thus pave the way for evaluation of a similar strategy in SPG11 patients. Such strategy could eventually be coupled with the use of immunomodulators that ameliorated the motor symptoms observed at a later stage in a preclinical study using the same animal model (Hörner et al., 2022).

## Acknowledgements

We thank the patients for their participation in research as well as the Phenoparc, iGenSeq, ICV, ICV-Vector, ICV-iPS, Histomics, ICM.quant and DNA and cell bank facilities of the Paris Brain Institute as well as the Chronobiotron of Strasbourg for their help. We also thank Stecy Chhor and Anne Robé for their help. This work was supported by“Investissements d’Avenir” program [ANR-10-IAIHU-06] and [ANR-11-INBS-0011] grants. The work was supported by funding from the European Research Council (ERC-POC grant No 825713 to F.D.), the Association Strumpell-Lorrain-HSP-France (to G.S.), the Agence Nationale de la Recherche (ANR-19-CE17-0022 to F.D and B.C), the Italian Ministry of Health (RC5X1000 to F.M.S) and the Tom Wahlig Stiftung (grant CC 02790-2020 to I.R.). D.S. received a fellowship from the French Ministry of Research (doctoral school ED3C). Lipidomics analyses by LC-HRMS/MS were funded and performed within the MetaboHUB French infrastructure (ANR-INBS-0010).

## Author contribution

G.S., S.C., B.Cow. and F.D. designed the study; M.F., M.C., S.B., J.Be., A.M., J.Br., I.R., S.M., K.D., K.H.E.H, B.Col., D.S., M.D. performed experimental work; K.D. provided reagents; M.F., M.C., S.B., J.Be., A.M., J.Br., I.R., S.M., K.D., B.Col., A.D., F.M.S., S.C, B.Cow. and F.D. analyzed data. S.C. and F.D. wrote the manuscript.

## Data availability statement

Data related to the current manuscript are available from F.D. upon reasonable request.

