## Supplementary Figures for "Decreasing ganglioside synthesis delays motor and cognitive symptom onset in *Spg11* knockout mice"

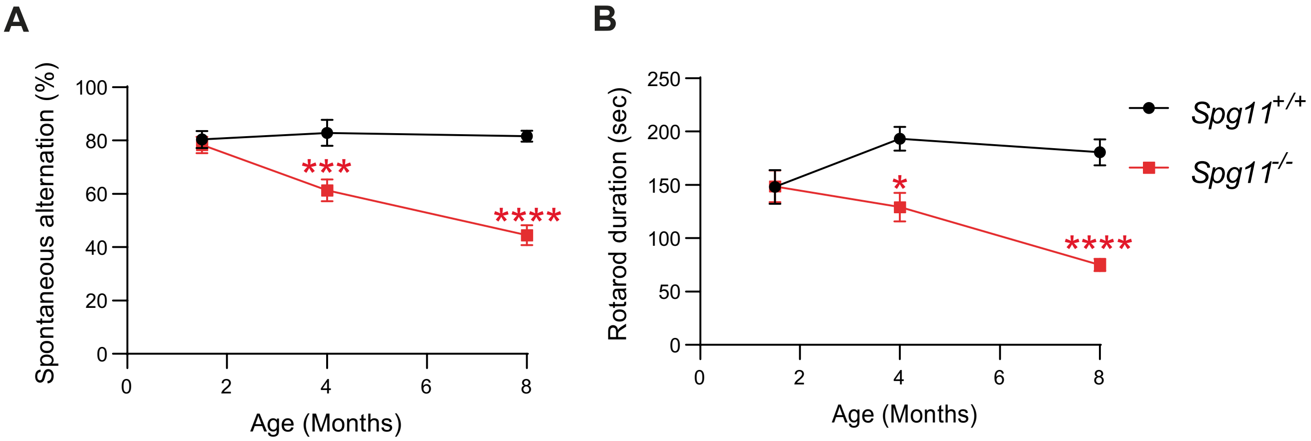


**Supplementary Figure 1. Natural history of cohort of *Spg11*$^{-/-}$ mice**. *Spg11*$^{+/+}$ and *Spg11*$^{-/-}$ mice housed in Chronobiotron facility (Strasbourg, France) were evaluated for the cognitive function using the Y-maze test (A) and the motor function using the accelerating rotarod test (B) at the ages of 1.5, 4 and 8 months. Mean and SEM. N>14 mice/group. *P<0.05; ***P<0.001; ****P<0.0001. Two-Way ANOVA followed by Holm-Sidak multiple comparison test.

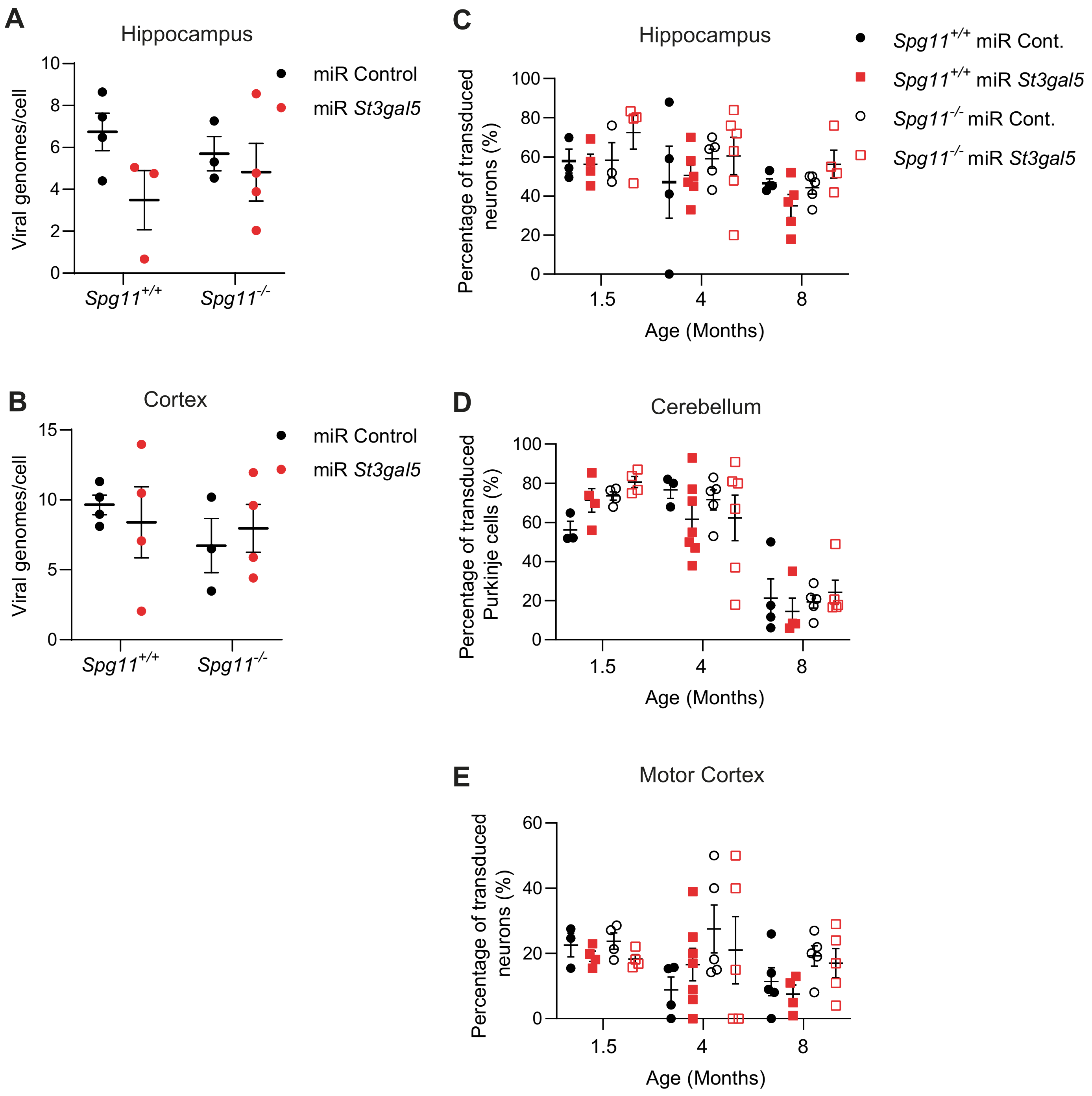


**Supplementary Figure 2. Transduction of neurons in various brain regions upon intravenous injection of AAV-PHP.eB.** *Spg11*$^{+/+}$ and *Spg11*$^{-/-}$ mice injected with AAV-PHP.eB expressing either a control miRNA or a miRNA downregulating *St3gal5* at the age of 3 weeks, were analyzed by qPCR at the age of 1.5 months (A,B) or immunohistochemistry at the age of 1.5, 4 or 8 months for the expression of GFP (C-E). The number of viral genome copies/cell was quantified in the hippocampus (A) and cortex (B). The proportion of neurons expressing GFP was monitored for each cohort in the CA3 region of the hippocampus (C), in Purkinje cells of the cerebellum (D) or in the motor cortex (E). Means and SEM. N=3-8 mice per group.

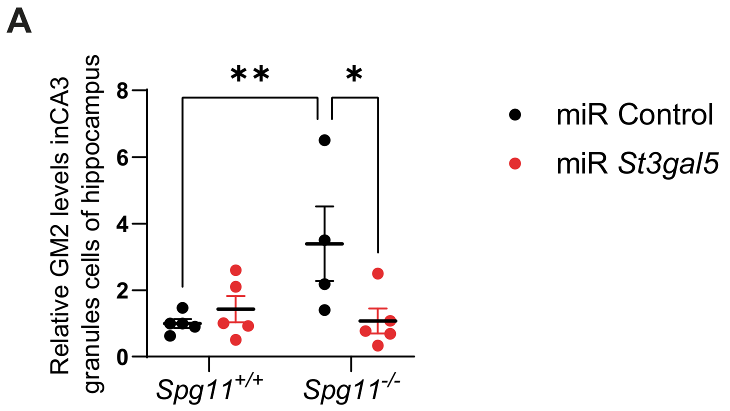


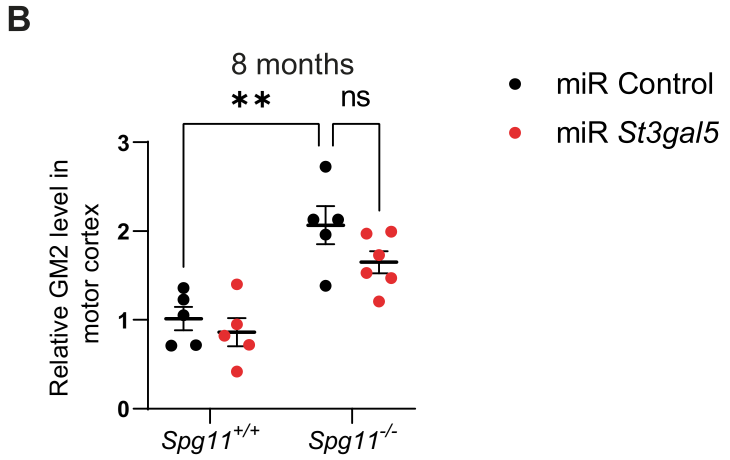


**Supplementary Figure 3. Downregulation of *St3gal5* delays accumulation of gangliosides in neurons of *Spg11*$^{-/-}$ mice.** (A) Quantification of the relative levels of GM2 ganglioside in neurons of the hippocampus of 4 month-old *Spg11*$^{+/+}$ and *Spg11*$^{-/-}$ mice injected with a control virus or AAV-PHP.eB downregulating *St3gal5*. (B) Quantification of the relative levels of GM2 ganglioside in neurons of the motor cortex of 8 month-old *Spg11*$^{+/+}$ and *Spg11*$^{-/-}$ mice injected with a control virus or AAV-PHP.eB downregulating *St3gal5*. Means and SEM. N=4-7 independent mice/group. *$P<0.05$; ** $P<0.01$; Two-way ANOVA followed by Holm-Sidak multiple comparison test.

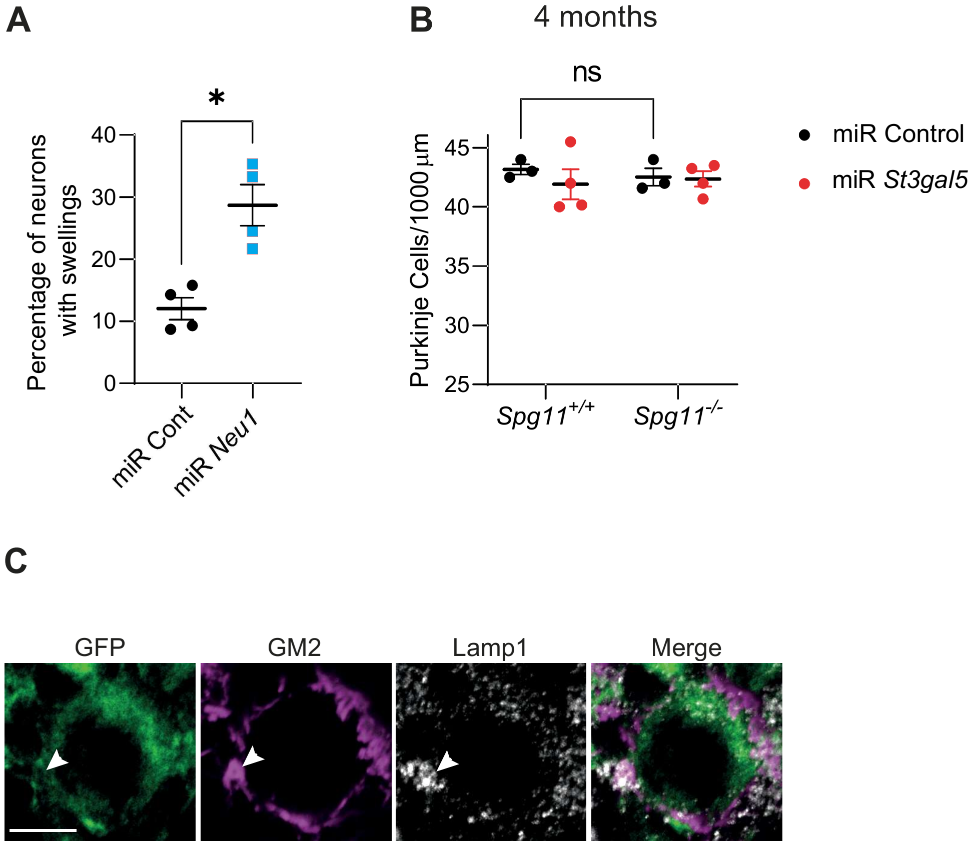


**Supplementary Figure 4. Ganglioside accumulation is associated with swellings in neurons**. (A) Quantification of the proportion of wild-type neurons presenting swellings when transfected with vectors expressing GFP and either a control miRNA or miRNA downregulating the neuroaminidase Neu1. Means and SEM. N= 4 independent experiments with more than 50 neurons analyzed per experiment. P<0.05; Mann Whitney test.
(B) Quantification of the number of Purkinje cells (measured in lobulus simplex) per 100 µm at the age of 4 months in *Spg11*$^{+/+}$ and *Spg11*$^{-/-}$ mice injected with AAV-PHP.eB expressing either a control miRNA or a miRNA downregulating *St3gal5*. Note the absence of neuronal death at that stage of the disease in *Spg11*$^{-/-}$ mice. N=3-5 independent mice/group. Two-way ANOVA followed by Holm-Sidak multiple comparison test. (C) Immunostaining of Purkinje cell of *Spg11*$^{-/-}$ mice injected with control AAV-PHP.eB immunostained with GFP, Lamp1 and GM2 antibodies. Note the accumulation of GM2 in swelling positive for Lamp1 (Arrowhead). Scale bar: 10 µm.
